# RPAP2 regulates a transcription initiation checkpoint by prohibiting assembly of preinitiation complex

**DOI:** 10.1101/2021.06.18.448918

**Authors:** Xizi Chen, Yilun Qi, Xinxin Wang, Zhenning Wang, Li Wang, Aixia Song, Jiabei Li, Dan Zhao, Hongwei Zhang, Qianwei Jin, Fei Xavier Chen, Yanhui Xu

**Author notes:** These authors contributed equally to this work (Xizi Chen, Yilun Qi, Xinxin Wang, Zhenning Wang, Li Wang, Aixia Song).

## Abstract

RNA polymerase II (Pol II)-mediated transcription in metazoan requires precise regulation. RNA polymerase II-associated protein 2 (RPAP2) was previously identified to transport Pol II from cytoplasm to nucleus and dephosphorylates Pol II C-terminal domain (CTD). We found that RPAP2 binds hypo/hyper-phosphorylated Pol II with undetectable phosphatase activity. Structure of RPAP2-Pol II shows mutually exclusive assembly of RPAP2-Pol II and pre-initiation complex (PIC) due to three steric clashes. RPAP2 prevents/disrupts Pol II-TFIIF interaction and impairs in vitro transcription initiation, suggesting a function in prohibiting PIC assembly. Loss of RPAP2 in cells leads to global accumulation of TFIIF and Pol II at promoters, indicating critical role of RPAP2 in inhibiting PIC assembly independent of its putative phosphatase activity. Our study indicates that RPAP2 functions as a gatekeeper to prohibit PIC assembly and transcription initiation and suggests a novel transcription checkpoint.

## Introduction

The sequence-specific transcription factors (TFs) bind regulatory elements (promoters and enhancers) and multi-subunit Mediator to facilitate the assembly of a preinitiation complex (PIC) and activate RNA polymerase II (Pol II)-mediated eukaryotic transcription initiation on target genes. PIC assembly involves sequential recruitment of general transcription factors (GTFs) TFIID, TFIIA, TFIIB, TFIIF-bound Pol II, TFIIE, and TFIIH (Roeder, 1996; Thomas and Chiang, 2006; Zawel and Reinberg, 1993). Within the 10-subunit TFIIH, the DNA translocase subunit (XPB) stimulates promoter opening (Guzder et al., 1994; Lin et al., 2005) and the cyclin-dependent kinase 7 (CDK7) phosphorylates Ser5 residues of the heptapeptide repeats (Y^1^S^2^P^3^T^4^S^5^P^6^S^7^) of the RPB1 C-terminal domain (CTD) (Feaver et al., 1994; Fisher and Morgan, 1994), the two processes required for transcription initiation. As a focal point of transcription regulation, activation of transcription initiation has been well-characterized through biochemical studies (Buratowski et al., 1989; Cortes et al., 1992; Flores et al., 1991) and was recently revealed by structural studies from our group and others (Abdella et al., 2021; Chen et al., 2021a; Chen et al., 2021b; Rengachari et al., 2021). Compared to extensive studies of positive regulation, negative regulation of transcription initiation is less understood.

The human RNA polymerase II-associated protein 2 (RPAP2) and its yeast homolog regulator of transcription protein (Rtr1) were originally identified to transport the associated Pol II from the cytoplasm to nucleus (Forget et al., 2013; Gibney et al., 2008). Cytoplasm accumulation of RPAP2 was observed in Myofibrillar myopathies, a group of neuromuscular disorders (Guglielmi et al., 2015). Subsequent studies showed that depletion of RPAP2/Rtr1 led to defect of transcription termination (Victorino et al., 2020) and increase in the level of Ser5-phosphorylation (pSer5) of Pol II CTD (Egloff et al., 2012; Hunter et al., 2016; Kim et al., 2009; Mosley et al., 2009), which serves as an indicator of enhanced transcription initiation. As RPAP2-binding partners, regulator of pre-mRNA-domain-containing 1A (RPRD1A) and RPRD1B form heterodimer and preferentially bind Pol II with CTD phosphorylated at Ser2/Ser7 (Ni et al., 2014) and acetylated at position 7 lysine residues (Ali et al., 2019), consistent with the interaction between hyperphosphorylated Pol II and Rtr1 (Smith-Kinnaman et al., 2014).

The role of RPAP2/Rtr1 in transcription regulation has been believed to be derived from the phosphatase activity of RPAP2 (Egloff et al., 2012) and Rtr1 (Mosley et al., 2009). However, in vitro biochemical analyses (Hsu et al., 2014; Xiang et al., 2012) indicated the RPAP2/Rtr1 possesses undetectable phosphatase activity or much lower activity compared to other known Pol II CTD phosphatases, such as Ssu72 (Zhang et al., 2011), Scp1 (Zhang et al., 2006), and Fcp1 (Hausmann and Shuman, 2002). Structural studies indicated that Rtr1 of Kluyveromyces lactis and Saccharomyces cerevisiae adopt similar fold, which lacks well-defined catalytic pocket of phosphatase (Hsu et al., 2014; Irani et al., 2016; Xiang et al., 2012). The lack of efficient phosphatase activity suggests that RPAP2/Rtr1 may involve transcription regulation through a phosphatase-independent function.

Here, we observed near stoichiometric association of endogenous RPAP2 during the purification of recombinant human Pol II. The in vitro assay showed undetectable phosphatase activity on phosphorylated Pol II CTD. The cryo-electron microscopy (EM) structures of RPAP2-Pol II complexes indicate mutually exclusive assembly of RPAP2-Pol II with PIC or elongation complex (EC). Biochemical analysis indicated that RPAP2 prevents Pol II-TFIIF interaction, disrupts Pol II-TFIIF complex, and inhibits in vitro transcription initiation. Chromatin immunoprecipitation (ChIP)-sequencing analysis showed that the deletion of RPAP2 led to an increase of TFIIF occupancy on promoters, indicative of enhanced assembly of PIC. Such negative effect on transcription initiation is independent of the putative phosphatase activity of RPAP2. Thus, we identified a transcription pre-initiation checkpoint, in which RPAP2 binds Pol II and prevents PIC assembly and transcription initiation.

## Results

### RPAP2 binds Pol II but shows undetectable phosphatase activity

During purification of human Pol II overexpressed in Expi293F cells, we observed a stably co-purified Pol II-binding partner (Figure 1A). Mass spectrometry (MS) analysis indicated that the protein is RNA polymerase II associated protein 2 (RPAP2), a previously identified Pol II-binding protein. We observed higher and lower bands of RPB1 in the purified Pol II, indicative of Pol II in the hyperphosphorylated (Pol IIo) and hypophosphorylated (Pol IIa) forms, respectively. Consistently, trace amount of phosphorylated Pol II (phosphorylation of CTD at Ser2, Ser5, and Ser7) could be detected from the purified RPAP2, which was overexpressed in Expi293F cells (Figure 2E, lane 1). To test whether RPAP2 binds Pol IIa or Pol IIo, we next separately purified RPAP2 and prepared phosphorylated Pol II as previously described (Zheng et al., 2020) (Figures 1B and 2E). The in vitro pulldown assay showed that RPAP2 binds Pol II in the two phosphorylation forms, indicating that Pol II CTD phosphorylation is not required for RPAP2-Pol II interaction.

**Figure 1.**
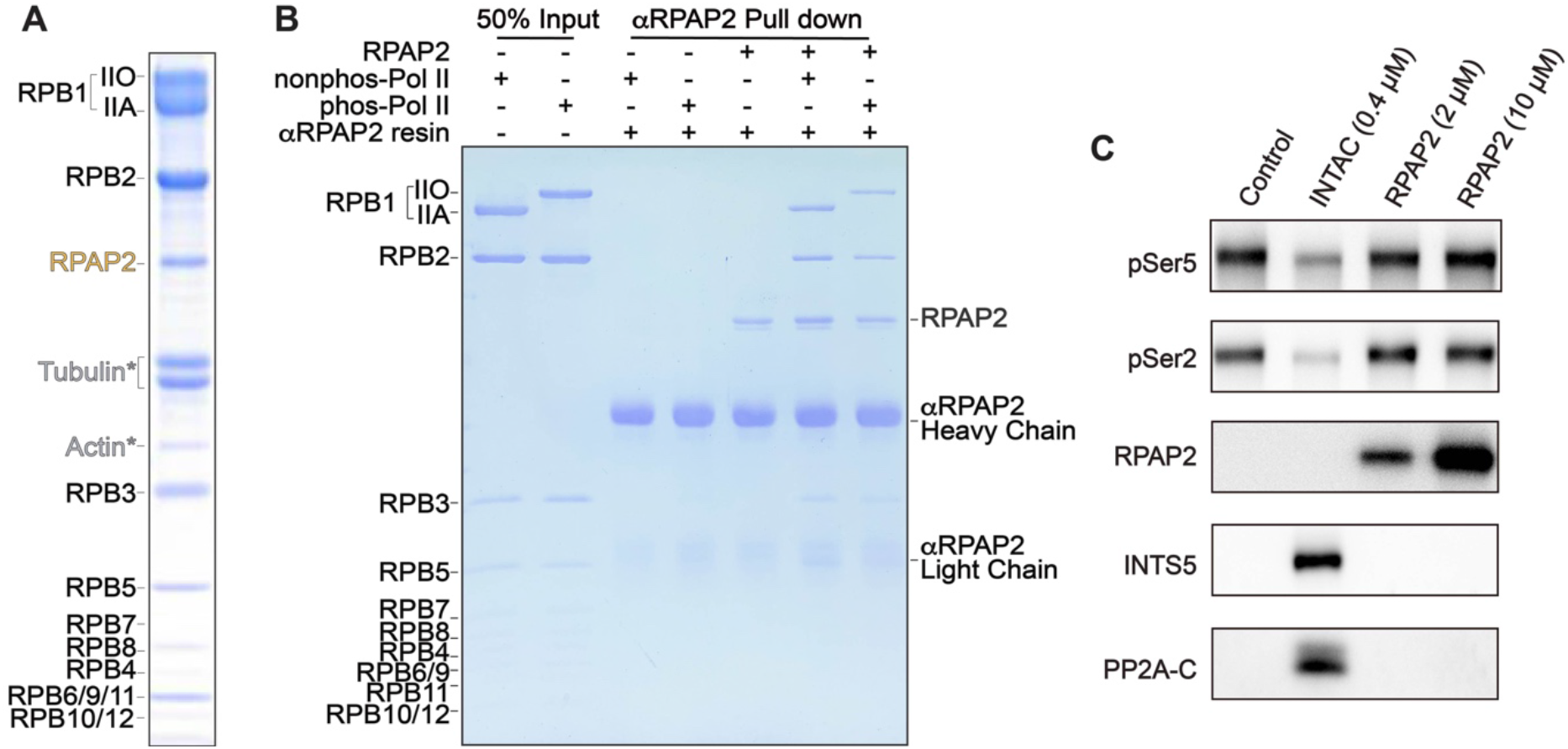
Purification and characterization of RPAP2-Pol II complex. (**A**) Endogenous RPAP2 was co-purified during purification of human Pol II that was overexpressed in Expi293F cells. The purified complex was subjected to SDS-PAGE followed by Coomassie blue staining. Contaminating proteins were indicated with stars. The higher and lower band represent RPB1 in Pol IIo and Pol IIa, respectively. (**B**) In vitro pulldown assay using unphosphorylated Pol II, phosphorylated Pol II and RPAP2. The bound proteins were subjected to SDS-PAGE followed by Coomassie blue staining. (**C**) In vitro phosphatase assay using purified RPAP2 and INTAC as enzymes and phosphorylated Pol II as substrate. The reactions were subjected to Western blotting using the indicated antibodies. The enzyme concentrations are indicated above each lane. See also Figure S1.

**Figure 2.**
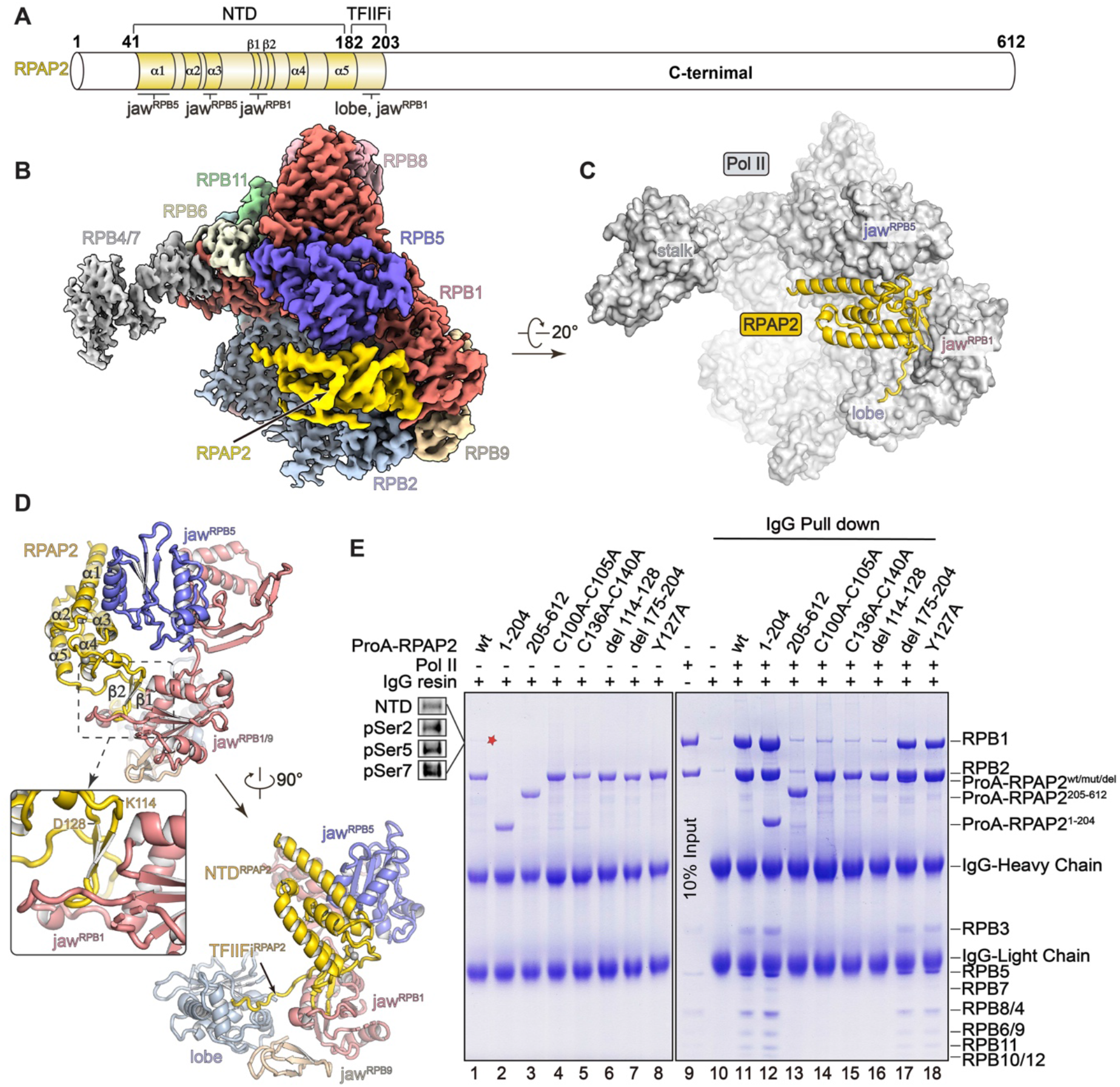
Overall structure of RPAP2-Pol II complex. (**A**) Domain structure of the RPAP2. The domains and regions built in structural model are colored and the unmodeled region is in white. Interfaces between RPAP2 and Pol II are indicated below. (**B**) A combined cryo-EM map of RPAP2-Pol II complex. (**C**) Structural model of RPAP2-Pol II complex with Pol II shown in surface and RPAP2 shown in cartoon. The zinc cation at the zinc finger of RPAP2 is shown as a gray ball. (**D**) Two views of intermolecular contacts between RPAP2 and Pol II. The interaction between the two-stranded β-sheet and RPB1 jaw is indicated with dashed box and shown in close-up view (middle panel). (**E**) Interactions between Pol II and RPAP2. The cell lysates containing various Protein A (ProA)-tagged RPAP2 mutants or truncations were applied to IgG resins, which were further incubated with purified Pol II. The unbound proteins were washed away and bound proteins were subjected to SDS-PAGE followed by Coomassie blue staining. ProA-tagged RPAP2 overlaps with RPB2 in lanes 11-18 and the amounts of ProA-RPAP2 are indicated in lanes 1-8. Note that trace amount of endogenous Pol II was pulled out by RPAP2^WT^ (lane 1). See also Figure S2, S3 and Table S1.

The human RPAP2 and its yeast homolog Rtr1 have been reported to possess protein phosphatase activity against phosphorylated Ser5 (pSer5) of Pol II CTD (Egloff et al., 2012; Hsu et al., 2014; Kim et al., 2009; Mosley et al., 2009). However, RPAP2-bound Pol II remained evidently hyperphosphorylated after the two-day purification (Figure 1A), suggesting inefficient dephosphorylation of RPAP2 on phosphorylated Pol II CTD.

To directly measure RPAP2 phosphatase activity, we performed an in vitro phosphatase assay using the phosphorylated Pol II as substrate, which possesses pSer5 and pSer2 of RPB1 CTD (Figure 1C, lane 1). As a positive control, integrator-containing PP2A complex (INTAC) at 0.4 μM concentration showed largely dephosphorylated RPB1 (lane 2), consistent with our previous study (Zheng et al., 2020). In contrast, RPAP2 at a concentration of as high as 10 μM showed undetectable phosphatase activity (lanes 3 and 4), suggesting that RPAP2 possesses very weak, if any, phosphatase activity against pSer5 of Pol II CTD. The result is consistent with previous structural and biochemical studies showing that that Rtr1 (RPAP2 homolog) lacks an active site and phosphatase activity (Xiang et al., 2012). Collectively, the in vitro assays suggest that RPAP2 does not efficiently dephosphorylates Pol II CTD and Pol II CTD phosphorylation is not required for binding of RPAP2 to Pol II, suggesting a phosphorylation independent role of RPAP2 in Pol II function.

### Structure of RPAP2-Pol II complex

We next determined the structure of human RPAP2-Pol II complex using cryo-EM single particle reconstruction and the cryo-EM map was refined to 3.5 Å resolution (Figures S1 and S2). RPRD1A-RPRD1B heterodimer was previously reported to bind RPAP2 and facilitate the recognition of phosphorylated Pol II CTD (Ni et al., 2014). We have also complexed *Sus scrofa* Pol II (four-residue substitution in human Pol II) with human RPAP2 and RPRD1A-RPRD1B followed by gradient fixation (Grafix) (Figure S1) (Kastner et al., 2008) and the cryo-EM map was refined to 2.8 Å resolution. The two cryo-EM maps showed almost identical conformation and RPRD1A-RPRD1B was not observed, consistent with the binding of RPRD1A-RPRD1B to the highly flexible Pol II CTD.

Structure determination was focused on RPAP2-Pol II-RPRD1A-RPRD1B (termed RPAP2-Pol II for simplicity) and the structure will be discussed below (Figure 2 and Video S1). The cryo-EM map around RPAP2 was locally refined to 3.4 Å resolution. For structural model building, the structural templates of Pol II from holo PIC (hPIC) complex (PDB: 7EGB) (Chen et al., 2021a) and yeast Rtr1 (PDB: 4FC8) (Xiang et al., 2012) were respectively docked into the cryo-EM map, followed by manual adjustment (Figure S2 and Table S1).

The modeled RPAP2 consists of an N-terminal domain (NTD^RPAP2^, residues 41-182) followed by an extended loop (residues 183-203), which we termed TFIIF inhibitory region (TFIIFi^RPAP2^, described below) (Figure 2A-2D and Video S1). The C-terminal region (204-612) was invisible in the cryo-EM map, consistent with the predicted flexibility. The NTD^RPAP2^ consists of a five-helix bundle and a characteristic zinc finger. The zinc finger and a short helix stabilize the five-helix bundle on two oppositive ends. The overall fold of NTD^RPAP2^ is generally similar to that of the reported structures of yeast Rtr1 (Hsu et al., 2014; Irani et al., 2016; Xiang et al., 2012) (Figure S3A).

In RPAP2-Pol II, Pol II adopts a similar conformation to that in PIC (Chen et al., 2021a; Chen et al., 2021b) and EC (Bernecky et al., 2016) (Figures 3A and S4A). The NTD^RPAP2^ is grasped by the RPB5 jaw and RPB1 jaw (Figure 2D). The five-helix bundle binds the parallel helices of the RPB5 jaw. A two-stranded β-sheet (residues 114-128) protrudes out of the five-helix bundle and packs against the RPB1 jaw. The tip (residues 120-125) of the β-sheet inserts into and stabilizes a flanking hairpin of RPB1 jaw, which was not previously modeled due to the lack of stabilization (Bernecky et al., 2016; Chen et al., 2021a; Chen et al., 2021b). The TFIIFi^RPAP2^ packs against the lobe of RPB2, which contacts the charge helix of TFIIF (TFIIFα subunit) in PIC/EC.

**Figure 3.**
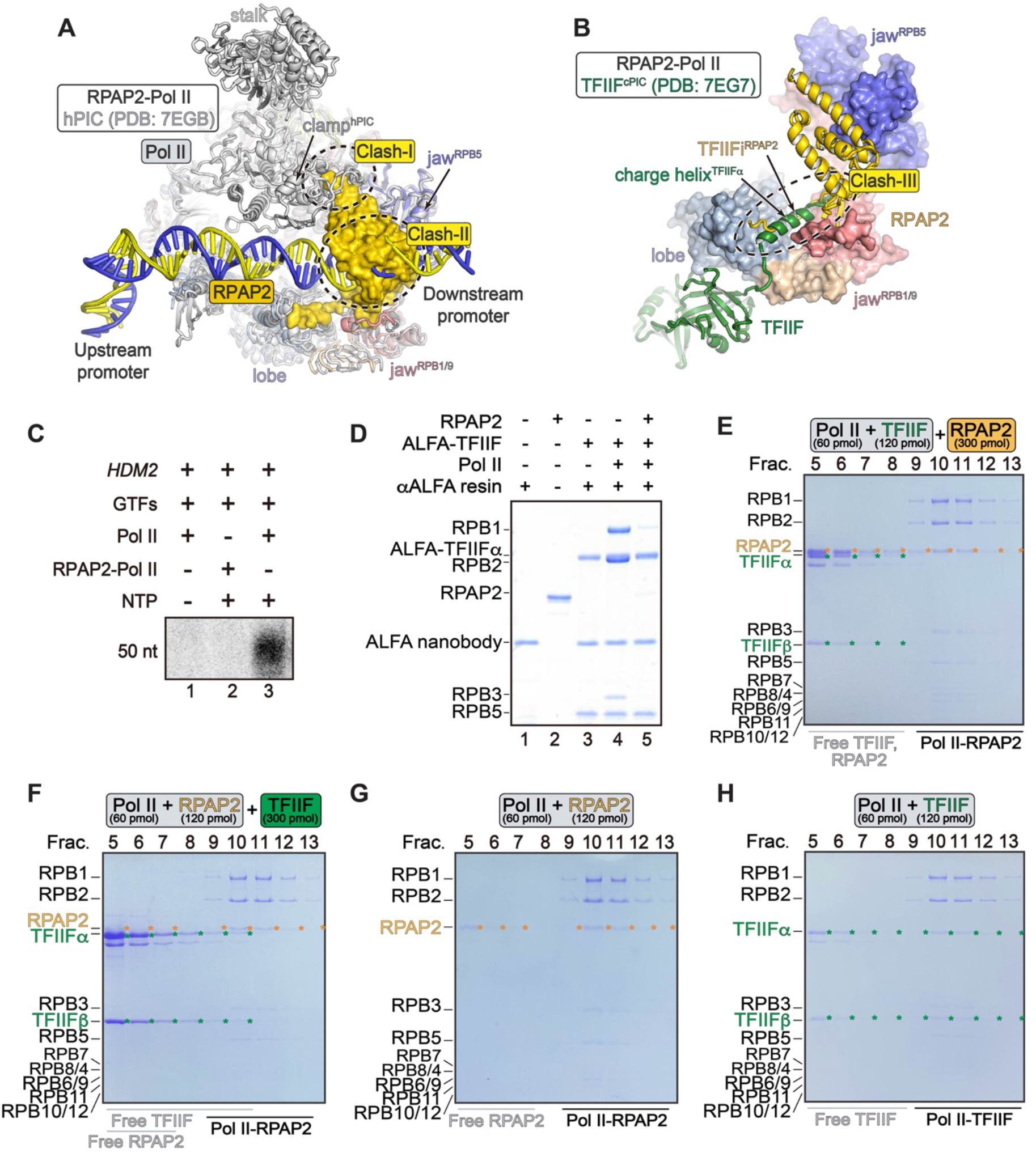
RPAP2 disrupts Pol II-TFIIF complex and inhibits PIC assembly. (**A**) Structures of RPAP2-Pol II and hPIC (PDB:7EGB) (Chen et al., 2021a) with Pol II superimposed. RPAP2-Pol II is colored as in Figure 2B and hPIC is colored in grey. The promoter is colored in yellow and blue for clarity. The GTFs of hPIC were omitted for complicity. RPAP2 is shown in surface representation. Steric clashes are indicated with dashed circle. (**B**) Structural comparison of RPAP2-Pol II with core PIC (cPIC) (PDB: 7EG7) (Chen et al., 2021a) with Pol II superimposed. Pol II is shown in surface and steric clash between TFIIF charge helix and TFIIiF^RPAP2^ is indicated with dashed circle. Unnecessary regions of cPIC were omitted for clarity. **(C)** Autoradiogram of in vitro transcription initiation reactions. The complexes in the reactions included purified GTFs (TFIIA, TFIIB, TFIID, TFIIF, TFIIE and TFIIH), *HDM2* promoter and either Pol II or RPAP2-Pol II as indicated. As previously described (Chen et al., 2021b), the reactions were incubated at 25°C for 30 min and then were subjected to urea polyacrylamide gels and autoradiography. **(D)** In vitro competitive binding assay using purified TFIIF (TFIIFα has an ALFA-tag), RPAP2 and Pol II. TFIIF and Pol II were pre-assembled and immobilized on anti-ALFA resins before adding an equal amount of RPAP2. The bound proteins were subjected to SDS-PAGE and stained using Coomassie blue. (**E-H**) RPAP2 prohibits and disrupts Pol II-TFIIF interaction. RPAP2 (300 pmol) was incubated with the pre-assembled Pol II-TFIIF (60 pmol) (E) or TFIIF (300 pmol) was incubated with the pre-assembled RPAP2-Pol II (60 pmol) (F), followed by glycerol density gradient ultracentrifugation. Fractions of glycerol density gradient centrifugation of RPAP2-Pol II (G) and TFIIF-Pol II (H) are shown as control. See also Figure S4.

### The N-terminal region of RPAP2 is necessary and sufficient for binding of Pol II

We next performed in vitro pulldown assay to test the interaction between RPAP2 and Pol II. Reciprocal pulldown assay indicated that full-length RPAP2 and the N-terminal region (residues 1-204) bound Pol II (Figures 2E, lanes 11-12 and S3A, lanes 18-19) whereas the C-terminal region (residues 205-612) showed undetectable binding of Pol II (Figure 2E, lane 13 and S3A, lane 20). Double-mutation C100A-C105A and C136A-C140A largely decreased the interaction (Figure 2E, lanes 14-15 and S3A, lane 21-22), indicating a critical role of the zinc-finger in maintaining the overall fold of NTD^RPAP2^. The deletion of the β-sheet (residues 114-128) decreased the interaction (Figures 2E, lanes 16 and S3A, lane 23), consistent with its position in bridging NTD^RPAP2^ and RPB1 jaw of Pol II.

It was reported that mutation Y105A of yeast Rtr1 (equivalent of Y127A in RPAP2) impairs phosphatase activity (Irani et al., 2016). The RPAP2-Pol II interaction was not obviously affected by the mutation Y127A (Figure 2E lane 18 and S3A, lane 25), suggesting a phosphatase-independent function of RPAP2 in binding of Pol II. Thus, RPAP2^Y127A^ represents a phosphatase-dead mutant that maintains RPAP2-Pol II interaction. Moreover, the deletion of extended loop region (residues 175-204) has no obvious effect on RPAP2-Pol II interaction.

### Three steric clashes between RPAP2 and PIC/EC elements

Comparison of RPAP2-Pol II structure with the structures of PIC (Chen et al., 2021a) and EC (Bernecky et al., 2016) complexes shows that the Pol II-bound RPAP2 may generate three steric clashes with structural elements of PIC or EC (Figures 3A, 3B and S4A). (1) The NTD^RPAP2^ generates steric clash (Clash-I) with the Pol II clamp in PIC/EC (Figures 3A and S4A), consistent with the absence of cryo-EM density of the clamp in RPAP2-Pol II reconstruction (Figure 2B). (2) The NTD^RPAP2^ generates an apparent clash (Clash-II) with DNA at the entry tunnel in PIC/EC, suggesting that binding of Pol II to RPAP2 and DNA are mutually exclusive. (3) Superimposition of RPAP2-Pol II and PIC shows obvious overlaps (Clash-III) of the TFIIFα charge helix and TFIIFi^RPAP2^ (Figure 3B and Video S2). The above structural analyses suggest that RPAP2 either inhibits PIC/EC assembly or dissociates from Pol II during assembly of PIC/EC if not undergoing significant conformational changes.

### RPAP2 disrupts Pol II-TFIIF interaction and prohibits PIC assembly

It is known that RPAP2 shuttles from the cytoplasm and nucleus with the associated Pol II (Forget et al., 2013; Gibney et al., 2008), suggesting that the Pol II-bound RPAP2 may function prior to the assembly of Pol II into PIC complex. To test whether RPAP2 affects transcription initiation, we performed in vitro transcription initiation assay using purified RPAP2, Pol II, and general transcription factors (GTFs) including TFIID, TFIIA, TFIIB, TFIIE, TFIIF, and TFIIH. As previously described (Chen et al., 2021b), Pol II and GTFs generated expected 50 nucleotide (nt) RNA products on *HDM2* promoter, indicating successful transcription initiation (Figure 3C, lane 3). In contrast, the pre-assembled RPAP2-Pol II showed undetectable transcription activity (Figure 3C, lane 2), indicating that RPAP2-bound Pol II does not support transcript initiation.

We next investigated in which step RPAP2 inhibits transcription initiation. It has been well-accepted that TFIIF is the first GTF that binds Pol II in step stepwise PIC assembly and that PIC is unable to be assembled in the absence of TFIIF (Cortes et al., 1992; Flores et al., 1991). Competitive pulldown assay showed that TFIIF stably binds Pol II and the Pol II-TFIIF complex is disrupted by the addition of RPAP2 (Figure 3D). Compared to standard Pol II-TFIIF assembly, much less TFIIF was observed in the peak fractions of Pol II in glycerol density gradient ultracentrifugation no matter whether TFIIF was added to the pre-assembled RPAP2-Pol II or RPAP2 was added to the pre-assembled Pol II-TFIIF (Figure 3E-3H). These results indicate that RPAP2 prohibits Pol II-TFIIF assembly and disrupts Pol II-TFIIF interaction but not vice versa. Consistently, an increasing amount of RPAP2 inhibits in vitro transcription initiation (Figure S4B).

We next performed a competitive binding assay to test whether other GTFs could disrupt RPAP2-Pol II interaction (Figure S4C-S4F). The immobilized RPAP2 was first incubated with Pol II followed by the addition of purified GTFs. The interaction between RPAP2 and Pol II was not obviously disrupted by TFIID-TFIIA-promoter, TFIIB, TFIIE, TFIIF, or TFIIH, in line with the architectural placement of RPAP2 on Pol II. The result is also consistent with the in vitro transcription assay (Figures 3C and S4B), suggesting that PIC assembly is inhibited by RPAP2 whereas RPAP2-Pol II interaction is not disrupted by GTFs in the in vitro system.

### RPAP2 regulates a pre-initiation checkpoint during cellular PIC assembly

To examine whether RPAP2 suppresses TFIIF recruitment and thus prohibits PIC assembly in cells, we depleted RPAP2 in human DLD-1 cells by shRNA (Figure 4A) and conducted ChIP with reference exogenous genome (ChIP-Rx) of TFIIF. The chromatin occupancy of TFIIF reflects dissociation of RPAP2 from Pol II and initial assembly of PIC complex on promoters. Strikingly, as shown by example genes (Figures 4B and S5A) and metagene analysis (Figure 4C), RPAP2 depletion greatly enhanced the binding of TFIIF at promoters, in supportive of the inhibitory role of RPAP2 in PIC assembly in cells. Consistently, the levels of Pol II increased at promoters upon RPAP2 depletion (Figure 4B and 4D). Measurement of total and elongating Pol II, represented by Pol II phosphorylated at Serine 2 of CTD, at gene bodies revealed an activation of transcription elongation, likely resulting from the increase in PIC assembly (Figure S5B-S5D).

**Figure 4.**
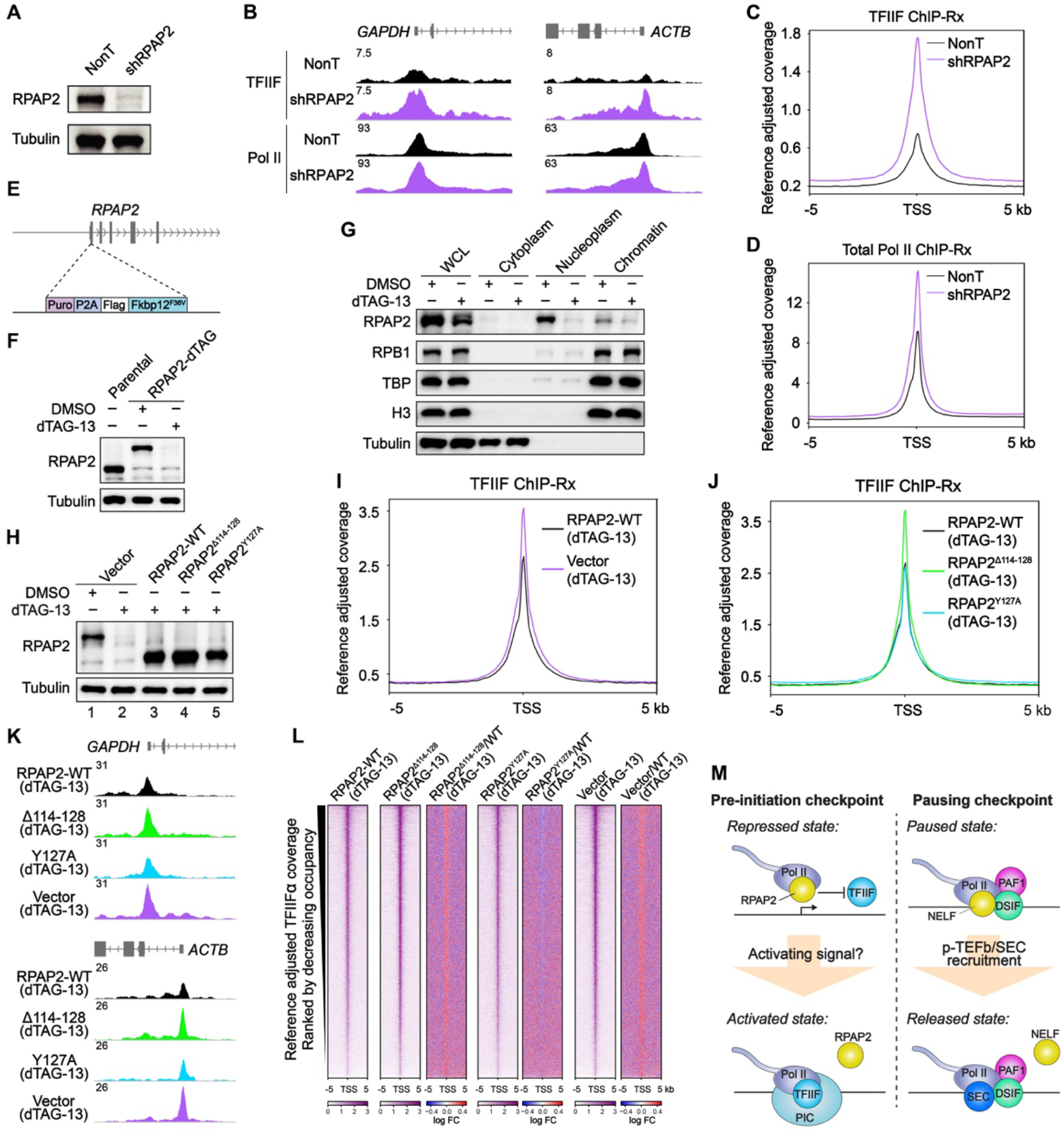
RPAP2 prohibits TFIIF recruitment and cellular PIC assembly independently of the putative phosphatase activity. **(A)** Western blots of whole cell extracts from DLD-1 cells with knockdown of RPAP2 by shRNA. **(B)** Representative track examples of genes *GAPDH* and *ACTB* showing the change of TFIIF and Pol II occupancy by RPAP2 knockdown. Pol II occupancy is represented by its largest subunit RPB1. (**C and D**) Metaplot showing the occupancy of TFIIF (C) and total Pol II (D) measured by ChIP-Rx in NonT and RPAP2 knockdown cells. **(E)** Schematic diagram of the generation of RPAP2-dTAG DLD-1 cells. **(F)** Western blots of whole cell extracts RPAP2-dTAG or parental cells treated with DMSO or dTAG for 3 hours. **(G)** Subcellular fractionation of RPAP2-dTAG treated with DMSO or dTAG followed by Western blotting. WCL, whole-cell lysate. **(H)** Induced expression of wildtype RPAP2, RPAP2^Δ114-128^, RPAP2^Y127A^, or vector in RPAP2-dTAG cells treated with dTAG-13, followed by Western of RPAP2. **(I)** Metaplot showing the occupancy of TFIIF in dTAG treated RPAP2-dTAG cells with induced expression of wildtype RPAP2 or vector. **(J)** Metaplot showing the occupancy of TFIIF in dTAG treated RPAP2-dTAG cells with induced expression of wildtype RPAP2, RPAP2^Δ114-128^, or RPAP2^Y127A^. **(K)** Representative track examples showing TFIIF occupancy in dTAG treated RPAP2-dTAG cells with induced expression of wildtype RPAP2, RPAP2^Δ114-128^, RPAP2Y127A, or vector. **(L)** Heatmaps of TFIIF occupancy centered at TSS of promoters ranked by decreasing occupancy. **(M)** Proposed model of the pre-initiation checkpoint regulated by RPAP2 (left). Pausing checkpoint (right) is shown for comparison. See discussion for detailed description.

To further confirm the direct regulation of PIC assembly by RPAP2, we utilized the degradation tag (dTAG) system (Nabet et al., 2018) by integrating the Flag-FKBP12^F36V^ tag at the N-terminus of the endogenous RPAP2 locus (RPAP2-dTAG) in DLD-1 cells (Figure 4E). The addition of dTAG-13 for three hours induced a rapid depletion of RPAP2 protein in RPAP2-dTAG cells (Figure 4F). Cellular fractionation showed a predominant presence of RPAP2 in nucleoplasm and much less RPAP2 on chromatin (Figure 4G, row 1), in line with our above results showing the mutually exclusive assembly of RPAP2-Pol II and PIC (Figure 3). Notably, rapid degradation of RPAP2 did not cause notably decreased association of Pol II on chromatin or accumulation of Pol II in cytoplasm (Figure 4G, row 2), the observed effect of siRNA-mediated RPAP2 silencing in previous study (Forget et al., 2013). This system allowed us to evaluate the direct role of RPAP2 in PIC assembly without affecting RPAP2 function in transporting Pol II.

To determine whether and, if yes, to what extent the Pol II binding capacity and putative phosphatase activity of RPAP2 contribute to its role in suppressing cellular PIC assembly, we generated rescue cell lines by respectively inducing the expression of wildtype and mutant RPAP2 in RPAP2-dTAG cells with rapid degradation of endogenous RPAP2 protein (Figure 4H, lanes 2-5). The two mutants include RPAP2^**Δ**114-128^, which compromises Pol II interaction (Figure 2E), and RPAP2^Y127A^, which represents a phosphatase-dead mutant (Figure 2E) (Irani et al., 2016). The comparison of TFIIF ChIP-Rx in dTAG-13 treated cells with induced expression of wildtype RPAP2 and an empty pLVX-Tet-On vector showed higher levels of TFIIF at promoters in the absence of RPAP2 (Figures 4I, 4K, and S5E, compare purple and black), revealing the direct role of RPAP2 in prohibiting PIC assembly. Despite being expressed to a higher level than wildtype RPAP2 (Figure 4H, lane 4), RPAP2^**Δ**114-128^ failed to rescue the aberrant accumulation of TFIIF (Figure 4, J and K, compare green and black). In contrast, the putative phosphatase-dead mutant RPAP2^Y127A^ fully restored the levels of TFIIF as the wildtype RPAP2 did (Figure 4J, 4K, and S5E, compare skyblue and black). Heatmaps showing the change of TFIIF levels indicated that wildtype RPAP2 and RPAP2^Y127A^, but not RPAP2^**Δ**114-128^, regulate TFIIF occupancy at genome-wide levels (Figure 4L). These results suggest that RPAP2 regulates a pre-initiation checkpoint through hindering the assembly of PIC and thus the following transcription activation. Importantly, this function of RPAP2 requires its association with Pol II but not the reported phosphatase activity.

## Discussion

Pol II-mediated transcription in metazoan is controlled at multiple levels including assembly of PIC on core promoters, transcription initiation, Pol II CTD phosphorylation and promoter escape, pausing and its release, elongation, and termination (Chen et al., 2018; Jonkers and Lis, 2015; Taatjes, 2021). Transcription checkpoints determine whether and when the transcription machinery pauses or proceeds to ensure transcription is under precise control. One of the most well-defined checkpoint is promoter-proximal Pol II pausing following the completion of initiation (Core and Adelman, 2019; Smith and Shilatifard, 2013) (Figure 4M, right panel). The generation and maintenance of paused Pol II rely on the coordination of several pausing factors including but not limited to the negative elongation factor (NELF), DRB sensitivity inducing factor (DSIF) and Pol II-associated factor 1 (PAF1) (Chen et al., 2018; Core and Adelman, 2019). Release of paused Pol II is driven by positive transcription elongation factor b (P-TEFb), a cyclin-dependent kinase 9 (CDK9)-containing complex, which phosphorylates Pol II and several key transcriptional regulators and thus activates the transcription machinery and relieves negative factors such as NELF.

In this study, we demonstrated that RPAP2 serves as a pivotal transcription gatekeeper in sterically inhibiting Pol II-TFIIF complex formation and transcription initiation. We proposed a transcription checkpoint prior to PIC assembly, which we termed pre-initiation checkpoint (Figure 4M, left panel). Given the critical role of TFIIF in stepwise PIC assembly (Cortes et al., 1992; Flores et al., 1991), an efficient transcription initiation requires the discharge of RPAP2 from Pol II before the formation and loading of Pol II-TFIIF at promoters. Our in vitro assays show that RPAP2 disrupts Pol II-TFIIF whereas TFIIF could not disrupt RPAP2-Pol II, suggesting additional factor(s) or post-translational modifications is required to disrupt RPAP2-Pol II and allow for Pol II-TFIIF formation.

The role of RPAP2/Rtr1 in transcription has mainly been attributed to its phosphatase activity towards pSer5 of Pol II (Ali et al., 2019; Egloff et al., 2012; Hsu et al., 2014; Hunter et al., 2016; Irani et al., 2016; Mosley et al., 2009; Ni et al., 2014; Victorino et al., 2020). However, this enzymatic activity is under debate due to the lack of a consensus phosphatase active pocket, as evidenced by our structural study and others (Irani et al., 2016; Xiang et al., 2012). Moreover, despite being reported as an atypical phosphatase, RPAP2/Rtr1 possesses very low in vitro phosphatase activity (Hsu et al., 2014; Irani et al., 2016). Consistently, our in vitro assays showed undetectable phosphatase activity of RPAP2 on phosphorylated Pol II CTD. Previous studies suggested that RPRD1A-RPRD1B heterodimer facilitates RPAP2 phosphatase activity by mediating RPAP2-Pol II association (Ali et al., 2019; Ni et al., 2014). However, we found that RPAP2 stably associates with Pol II (Pol IIa and Pol IIo) in the absence of RPRD1A-RPRD1B and that RPRD1A-RPRD1B leads to no apparent impact on the conformation of RPAP2-Pol II. In addition, our ChIP-Rx analyses showed that the phosphatase-dead mutant of RPAP2 fully rescues the aberrant accumulation of TFIIF caused by RPAP2 loss, indicating that RPAP2 modulates cellular PIC assembly independently of its putative phosphatase activity. Regarding its function as a specific pSer5 phosphatase, we surmise that the induced phosphorylation of Pol II upon RPAP2 depletion might partially result from effects of enhanced PIC assembly and thus transcriptional activation. Supporting this notion, our results showed that RPAP2 loss leads to a global increase in the occupancy of pSer2, which is not the substrate of RPAP2 but will be induced by transcriptional activation.

The function of RPAP2 in disrupting Pol II-TFIIF interaction is reminiscent of Gdown1, which also inhibits the binding of TFIIF to Pol II (Cheng et al., 2012; Jishage et al., 2012; Wu et al., 2012), consistent with the critical role of TFIIF in Pol II-mediated transcription. However, RPAP2 and Gdown1 regulate transcription in distinct stages. Gdown1 does not inhibit PIC assembly or transcription initiation (Mullen Davis et al., 2014), but instead, negatively regulates transcription at post-initiation stages (Cheng et al., 2012; DeLaney and Luse, 2016; Guo et al., 2014; Jishage et al., 2012; Jishage et al., 2018). Reconciling these available lines of evidence, we envision that activating signal-induced dissociation of RPAP2 and association of TFIIF occurs before PIC assembly; Gdown1 then competes with TFIIF for Pol II binding to hinder transcription at either pausing or elongation stages, given reported roles of TFIIF and Gdown1 in both pausing and transcription elongation (Cheng et al., 2012; DeLaney and Luse, 2016; Espinosa, 2012; Joo et al., 2019; Schweikhard et al., 2014).

During the preparation of this manuscript, a structure of RPAP2-Pol II complex was reported (Fianu et al., 2021), showing the incompatibility between RPAP2 and DNA in binding Pol II. The structure is generally similar to our RPAP2-Pol II structure (Figure S3B). Intriguingly, gel filtration analysis showed that the N-terminal region (1-215) of RPAP2 can be displaced from Pol II during PIC assembly in a TATA box-binding protein (TBP)-based system (Fianu et al., 2021). In contrast, we found that full-length RPAP2 is hardily displaced from Pol II and prohibits Pol II-TFIIF interaction during PIC assembly in TFIID-based system. Consistently, rapid degradation of RPAP2 leads to an aberrant accumulation of TFIIF at promoters with unnoticeable defects in Pol II biogenesis and nuclear import. The above distinct observations may result from differences in compositions of RPAP2 and PIC.

## Acknowledgments

We thank the Center of Cryo-Electron Microscopy of Fudan University for the supports on data collection and the Biomedical Core Facility, Fudan University for the support on mass spectrometry analyses. This work was supported by grants from the National key R&D program of China (2016YFA0500700), the National Natural Science Foundation of China (32030055, 31830107, 31821002), the Shanghai Municipal Science and Technology Major Project (2017SHZDZX01), Shanghai Municipal Science and Technology Commission (19JC1411500), the National Ten-Thousand Talent Program (Y. X.), the National Program for support of Top-Notch Young Professionals (Y. X.), and the Strategic Priority Research Program of the Chinese Academy of Sciences (grant no. XDB08000000).

## Author contributions

C. X. and X. W. prepared the samples for structural and biochemical analyses with the help from J. L., D. Z. and H. Z.; Y. Q. performed the cryo-EM analyses and the model building; Z. W. and L. W. generated the knock-in cell lines and conducted sequencing experiments with the help from Q.J.; A. S. analyzed the sequencing data; Y. X., F.X.C., and X. C. wrote the manuscript; Y. X. supervised the project.

## Data and materials availability

The cryo-EM maps have been deposited in the Electron Microscopy Data Bank (EMDB) with accession numbers of 31450 and 31451 and the structure coordinate has been deposited in the Protein Data Bank (PDB) with the accession number of 7F4G.

## Materials and Methods

### Antibodies and cell culture

Antibodies were as follows: Pol II (NTD) (#14958, Cell Signaling), Pol II (pSer2) (#13499, Cell Signaling), GTF2F1 (10093-2-AP, Proteintech), FLAG (#SLAB01, Smart Lifesciences), histone H3 (#4499, Cell Signaling), TBP: (66166-1-Ig, Proteintech). RPAP2 antibody is generated by Abclonal. 293T, DLD-1 and MEF cells were grown in DMEM supplemented with 10% FBS.

### Protein expression and purification

The 12 full-length open reading frames (ORFs) of human Pol II subunits were amplified from 293T cDNA by PCR or synthesized and sub-cloned into a modified pCAG vector and RPB1 was tagged with a C-terminal Protein A. All plasmids were co-transfected to Expi293F cells using PEI. After cultured at 37°C for 48 h, cells were harvested and lysed in buffer containing 30 mM HEPES pH 8.0, 300 mM NaCl, 0.25% CHAPS, 5 mM ATP, 5 mM MgCl_2_, 1 mM EDTA, 10 μM ZnCl_2_, 3 mM DTT, 10% Glycerol (v/v), 1 mM PMSF, 1 μg/mL Aprotinin, 1 μg/mL Pepstatin, 1 μg/mL Leupeptin at 4°C for 30 min. The cell lysate was clarified by centrifugation at 15,000 rpm at 4°C for 30 min and the supernatant was incubated with IgG resin (Smart-Lifesciences) for 1.5 h followed by on-column digestion at 4°C overnight in buffer containing 30 mM HEPES pH 8.0, 300 mM NaCl, 0.1% CHAPS, 1 mM EDTA, 2 mM MgCl_2_, 10 μM ZnCl_2_, 3 mM DTT, 10% Glycerol (v/v). The immobilized proteins were eluted out and further purified by a mono Q 5/5 column (GE Healthcare) and a Superose S6 Increase 5/150 GL column (GE Healthcare) in buffer containing 30 mM HEPES pH 8.0, 150 mM NaCl, 2 mM MgCl_2_, 0.2 mM EDTA, 10 μM ZnCl_2_ and 1mM TCEP. The peak fractions corresponding to the RPAP2-Pol II complex were concentrated using a 100-kDa cut-off centrifugation filter unit (Amicon Ultra). The purified complex was used for cryo-EM.

Human RPAP2 was sub-cloned into a modified pCAG containing an N-terminal protein A tag. The plasmid was transfected to Expi293F cells using PEI. After cultured at 37°C for 72 h, cells were harvested and lysed in Lysis buffer containing 30 mM HEPES pH 8.0, 300 mM NaCl, 0.25% CHAPS, 5 mM ATP, 5 mM MgCl_2_, 1mM EDTA, 10 μM ZnCl_2_, 3 mM DTT, 10% Glycerol (v/v), 1 mM PMSF, 1 μg/mL Aprotinin, 1 μg/mL Pepstatin, 1 μg/mL Leupeptin at 4°C for 30 min. Clarified lysates were applied to IgG resin (Smart-Lifesciences) for 1.5 h followed by on-column digestion at 4°C for 1 h. The immobilized proteins were eluted and further purified by a mono Q 5/5 column (GE Healthcare) and a Superdex75 10/300 GL column (GE Healthcare) in a buffer containing 30 mM HEPES pH 7.9, 100 mM KCl, 10 μM ZnCl_2_, 1 mM TCEP, 5% Glycerol (v/v). The peak fractions were pooled, aliquoted, snap frozen and stored at −80°C.

RPRD1A-RPRD1B complex was prepared essentially in a similar scheme. The two full-length ORFs of human RPRD1A and RPRD1B were separately sub-cloned into a modified pCAG vector containing N-terminal Protein A (ProA) tag and co-expressed in Expi293F cells. After cell lysis, the lysate was applied onto the IgG (Smart-Lifesciences) affinity chromatography column, followed by on-column digestion and the eluate was further purified by a mono Q 5/5 column (GE Healthcare) and a Superdex75 10/300 GL column (GE Healthcare) in buffer containing 30 mM HEPES pH 7.9, 100 mM KCl, 1 mM TCEP, 5% Glycerol (v/v). The peak fractions containing RPRD1A-RPRD1B complex were pooled, aliquoted, snap frozen and stored at −80°C.

GTFs (TFIID, TFIIA, TFIIB, TFIIF, TFIIE, TFIIH), *HDM2* promoter DNA, TFIID-TFIIA-promoter were prepared as previously described (Chen et al., 2021a). Unphosphorylated Pol II was isolated from *S. scrofa* thymus and purified following previously established protocol (Vos et al., 2018). Four residue substitutions (G882S of RBP2, T75I of RPB3, S140N of RPB3, and S126T of RPB6) exist between *S. scrofa* and *H. sapiens* Pol II. The phosphorylated Pol II was prepared by in vitro phosphorylation using TFIIH as previously described (Zheng et al., 2020).

### In vitro pull-down assay

In the RPAP2 pulldown assay, the purified RPAP2 was incubated with phosphorylated Pol II or unphosphorylated Pol II at 4°C for 2h in 400 μl Binding buffer containing 30 mM HEPES-KOH pH 7.9, 100 mM KCl, 0.05% CHAPS, 2mM MgCl_2_, 2 mM DTT, 0.2 mM EDTA, 10 μM ZnCl_2_ and 5% (v/v) glycerol. RPAP2 antibody (αRPAP2) and protein G resins were incubated at 4°C for 2h. Subsequently, the resins were washed three times and added to the RPAP2 and phosphorylated Pol II/ unphosphorylated Pol II mixture for another 2 hours at 4° C. The resins were extensively washed with the Binding buffer, and the bound proteins were subjected to SDS-PAGE followed by Coomassie blue staining.

In the 8WG16 pulldown assay, the purified RPAP2 mutants/truncations were individually incubated with purified Pol II at 4°C for 2h in Binding buffer, followed by the addition of Pol II antibody (8WG16) and protein G resins and further incubation at 4 ° C for 2h. After extensively washed using Binding buffer, the bound proteins were subjected to SDS-PAGE followed by Coomassie blue staining.

In the IgG pulldown assay, the indicated plasmids of N-terminal Protein A tagged RPAP2 mutants and truncations were individually transfected to Expi293F cells and cultured at 37°C for 72h. Cells of each RPAP2 protein were separately lysed as described above. The supernatant of cell lysate was incubated with IgG resins at 4°C for 2h. The resins were washed three times and resuspended 400 μl Binding buffer. The purified unphosphorylated Pol II was then added and incubated with the resins at 4°C for 2h. The resins were extensively washed and the bound proteins were subjected to SDS-PAGE and stained by Coomassie blue.

### In vitro phosphatase assay

In vitro phosphatase assay was performed as previously described (Zheng et al., 2020). Briefly, the phosphorylated Pol II (0.1 μM) was incubated with RPAP2 (2 μM, 10 μM) or INTAC complex (0.4 μM) in a final volume of 20 μl containing 50 mM HEPES-NaOH pH 7.4, 100 mM NaCl, 0.01% CHAPS, 10mM MgCl_2_, 1mM MnCl_2_, and 2 mM DTT. The reactions were performed at 30°C for 30 min and stopped by adding 5 μl of 5 × SDS loading buffer. Samples (2.5 μl) were subjected to SDS-PAGE and analyzed by Western blotting with indicated antibodies.

### In vitro competitive binding Assay

The competitive binding assay for RPAP2 and Pol II-TFIIF. The plasmids of N-terminal ALFA-tagged (Gotzke et al., 2019) TFIIF (ALFA-tagged TFIIFα and untagged TFIIFβ) were co-transfected to Expi293F cells and cultured at 37°C for 72h. Cells were pelleted and lysed as described above. The supernatant of cell lysate was incubated with ALFA-nanobody (αALFA) coupled resin at 4°C for 2h. The resins were washed three times and resuspended in Binding buffer. Subsequently, the purified unphosphorylated Pol II was added and incubated with the resins at 4°C for 2h. The resins were washed three times to remove unbound Pol II, followed by the addition of an equal amount of RPAP2 and further incubated for 2 hours. The resins were extensively washed and subjected to SDS-PAGE and stained by Coomassie blue.

The competitive binding assay for PIC components (TFIID-TFIIA-promoter, TFIIB, TFIIF, TFIIE and TFIIH) and RPAP2-Pol II. The purified RPAP2 was incubated with unphosphorylated Pol II at 4°C for 2h in 400 μl Binding buffer. RPAP2 antibody (αRPAP2) and protein G resins were incubated at 4°C for 2h. Subsequently, the resins were washed three times and added to pre-assembled RPAP2-Pol II for another 2 hours at 4°C. The resins were washed three times to remove unbound Pol II, followed by the addition of three molar excess of TFIID-TFIIA-promoter, TFIIB, TFIIF, TFIIE or TFIIH and further incubated for 2 hours. 10 μM THZ1 (MedChemExpress) was added to the reaction containing TFIIH to impaired the kinase activity of TFIIH. The resins were extensively washed with the Binding buffer, and the bound proteins were subjected to SDS-PAGE followed by Coomassie blue staining.

### Glycerol density gradient ultracentrifugation and competitive assay

To assemble the RPAP2-Pol II complex, 60 pmol of the purified *S. scrofa* unphosphorylated Pol II was incubated with 2-fold molar excess of purified RPAP2 at 4°C for 3h, and then added on top of a 4 ml 10%-50% (w/v) glycerol gradient in buffer containing 30 mM HEPES-KOH pH 7.9, 100 mM KCl, 2 mM MgCl_2_, 2 mM DTT and centrifuged for 16 h at 34,000 rpm at 4°C for 16 h using an SW60 Ti rotor (Beckman Coulter). The fractions, 200μl each, were subjected to SDS-PAGE and Coomassie blue staining. The peak fractions containing Pol II-RPAP2 complex were pooled, concentrated, aliquoted, snap frozen and stored at −80°C. The assembly of Pol II-TFIIF complex was prepared and analysis in a similar scheme.

To test the binding of RPAP2 and Pol II in the presence of excess TFIIF, 60 pmol of the purified *S. scrofa* unphosphorylated Pol II was pre-incubated with 2-fold molar excess of purified RPAP2 at 4°C for 1 h, followed by the addition of 5-fold molar excess of TFIIF and further incubation at 4°C for 2 h. The sample was subjected to glycerol density gradient ultracentrifugation as above mentioned.

The test of the binding of TFIIF and Pol II in the presence of excess RPAP2 were performed in a similar way. 5-fold molar excess of RPAP2 was incubated with 60 pmol of pre-incubated Pol II-TFIIF, and then analysis by glycerol density gradient ultracentrifugation.

To test the assembly of RPAP2-Pol II-RPRD1A-RPRD1B complex, The purified *S. scrofa* Pol II was incubated with RPAP2 and RPRD1A-PRD1B in molar ratio of 1:2:5 at 4°C for 3h, followed by glycerol density gradient ultracentrifugation in buffer containing 10%-50% (w/v) glycerol, 30 mM HEPES-KOH pH 7.9, 100 mM NaCl, 2 mM MgCl_2_, 10 μM ZnCl_2_, 2 mM DTT and centrifuged for 16 h at 34,000 rpm at 4°C for 16 h using an SW60 Ti rotor (Beckman Coulter). The fractions were subjected to SDS-PAGE and Coomassie blue staining.

### Cryo-EM sample preparation

To assemble the RPAP2-Pol II-RPRD1A-RPRD1B complex for cryo-EM sample preparation. The purified *S. scrofa* Pol II was incubated with RPAP2 and RPRD1A-PRD1B in molar ratio of 1:2:5 at 4°C for 3h. The mixture was then subjected to GraFix (Kastner et al., 2008). The glycerol gradient was prepared using light buffer containing 10% (w/v) glycerol, 30 mM HEPES-KOH pH 7.9, 100 mM NaCl, 2 mM MgCl_2_, 10 μM ZnCl_2_, 2 mM DTT, and heavy buffer containing 50% (w/v) glycerol, 30 mM HEPES-KOH pH 7.9, 100 mM NaCl, 2 mM MgCl_2_, 10 μM ZnCl_2_, 2 mM DTT, and 0.005% glutaraldehyde. The samples were centrifuged at 34,000 rpm at 4°C for 16 h using an SW60 Ti rotor (Beckman Coulter). Subsequently, fractions containing cross-linked complexes were quenched with 50 mM Tris pH 7.4 (25 °C). The homogeneity of peak fractions was assessed by negative stain EM. Fractions of interest were pooled, concentrated, followed by buffer exchange into a buffer containing 30 mM HEPES-KOH pH 7.9, 100 mM NaCl, 2 mM MgCl_2_, 10 μM ZnCl_2_, 2mM DTT, and 0.8 % (v/v) glycerol.

For negative-stain EM, 5 µl of freshly purified protein sample was applied onto a glow-discharged copper grid supported by a continuous thin layer of carbon film for 60 s before negative staining by 2% (w/v) uranyl formate at room temperature. The negatively stained grid was loaded onto a FEI Talos L120C microscope operated at 120 kV, equipped with a Ceta CCD camera.

For cryo-EM grid preparation, 4 μl of protein sample (about 1.5 mg/ml) was applied onto a glow-discharged holey carbon grid (Quantifoil Au, R1.2/1.3, 300 mesh). At a temperature of 4°C and under a humidity of 100%, the grid was blotted for 6 s using a FEI Vitrobot Mark IV and plunge frozen in liquid ethane cooled by liquid nitrogen. The grids were prepared in the H_2_/O_2_ mixture for 20 s using a Gatan 950 Solarus plasma cleaning system with a power of 5 W.

### Data collection

The cryo-EM grids of human RPAP2-Pol II and RPAP2-Pol II-RPRD1A-RPRD1B were loaded onto a Thermo Fisher Scientific Titan Krios transmission electron microscope equipped with a Gatan K2 direct electron detector and operating at 300kV for data collection. For RPAP2-Pol II, all the cryo-EM images were automatically recorded in the super-resolution counting mode using Serial-EM (Mastronarde, 2005) with a nominal magnification of 105,000x, which yielded a super-resolution pixel size of 0.678 Å, and with a defocus ranged from 1.8 to 2.5 μm. For RPAP2-Pol II-RPRD1A-RPRD1B, all the cryo-EM images were automatically recorded in the super-resolution counting mode using Serial-EM with a nominal magnification of 130,000x, which yielded a super-resolution pixel size of 0.522 Å, and with a defocus ranged from 1.8 to 2.5 μm. Each micrograph stack was dose-fractionated to 32 frames with a total electron dose of ∼ 50 e^-^/ Å^2^. 1,505 micrographs of RPAP2-Pol II and 2,529 micrographs of RPAP2-Pol II-RPRD1A-RPRD1B were collected for further processing.

### Image processing

For cryo-EM data, drift and beam-induced motion correction were applied on the super resolution movie stacks using MotionCor2(Zheng et al., 2017) and binned twofold to a calibrated pixel size of 1.356 Å/pixel and 1.044 Å/pixel, respectively. The defocus values were estimated by Gctf (Zhang, 2016) from summed images without dose weighting. Other procedures of cryo-EM data processing were performed using RELION 3.0(Zivanov et al., 2018) and cryoSPARC using dose-weighted micrographs.

For human RPAP2-Pol II, a subset of ∼10,000 particles were picked by RELION 3.0 without reference and subjected to reference-free 2D classification. Some of the resulting 2D class averages were low-pass filtered to 20 Å and used as references for automatic particle picking of the whole datasets in RELION resulting in an initial set of 1,593,921 particles for reference-free 2D classification. 862,194 particles were selected after several rounds 3D classifications, using a 60 Å low-pass filtered initial model from our previous cryo-EM reconstruction. 543,699 particles in four of the selected eight classes were imported to a 3D auto-refine and Postprocess, yielding a reconstruction of the RPAP2-Pol II complex at 3.5 Å resolution. For the RPAP2 module, two rounds of local mask 3D classification were performed. 85,919 particles in one of the selected three classes were imported to a 3D auto-refine and Postprocess, yielding a 4.5 Å reconstruction.

For RPAP2-Pol II-RPRD1A-RPRD1B, a subset of ∼10,000 particles were picked by RELION 3.0 without reference and subjected to reference-free 2D classification. Some of the resulting 2D class averages were low-pass filtered to 20 Å and used as references for automatic particle picking of the whole datasets in RELION resulting in an initial set of 1,173,686 particles for reference-free 2D classification. 754,747 particles were selected after several rounds of 3D classifications, using a 60 Å low-pass filtered initial model from our previous cryo-EM reconstruction. 646,517 particles in one of the selected six classes were imported to a cryoSPARC package for NU-refinement and sharpening, yielding a reconstruction of the RPAP2-Pol II-RPRD1-RPRD1B at 2.8 Å resolution. For the RPAP2 module, a local mask 3D classification were performed. 164,816 particles in one of the selected eight classes were imported to a cryoSPARC package for local refinement and sharpening, yielding a 3.4 Å reconstruction.

All reported resolutions were calculated based on the gold-standard Fourier shell correlation (FSC)=0.143 criterion. The GSFSC curves were corrected for the effects of a soft mask with high-resolution noise substitution. All cryo-EM maps were sharpened by applying a negative B-factor estimated in RELION. All the visualization and evaluation of the 3D volume map were performed within UCSF Chimera (Pettersen et al., 2004) or UCSF ChimeraX (Goddard et al., 2018), and the local resolution variations were calculated using RELION.

### Model building and structure refinement

The cryo-EM maps of RPAP2-Pol II-RPRD1A-RPRD1B (2.8 Å) and locally refined maps of RPAP2 (3.4 Å) were used for model building.

The following structural templates were used as references for model building. The structural templates include the cryo-EM structure of Pol II (PDB: 7EGB) (Chen et al., 2021a) and yeast Rtr1 (PDB: 4FC8) (Xiang et al., 2012). These structural models were docked into corresponding cryo-EM maps, followed by rigid-body fitting using UCSF Chimera(Pettersen et al., 2004) and manual adjustment in COOT(Emsley and Cowtan, 2004). Atomic structural models of Pol II and RPAP2 were built according to cryo-EM maps and refined in real space using Phenix(Adams et al., 2010).

Statistics of the map reconstruction and model refinement are shown in Table S1. The final models were evaluated using MolProbity (Chen et al., 2010). Model representations in the figures and movies were prepared by PyMOL (DeLano, 2002) or UCSF ChimeraX (Goddard et al., 2018).

### In vitro transcription initiation assay

GTFs and *HDM2* promotor DNA were prepared as previously described (Chen et al., 2021a). In vitro transcription initiation assay was performed as previously described (Cevher et al., 2014; Fujiwara and Murakami, 2019). Briefly, 1.3 pmol of *HDM2* promoter DNA was combined with 1.5 pmol of TFIID, 3 pmol of TFIIA, 3 pmol of TFIIB, 3 pmol of TFIIF, 3 pmol of TFIIE, 1.5 pmol of TFIIH, either 2 pmol of *S. scrofa* Pol II with increasing amount of RPAP2 (0 pmol, 2 pmol, 4 pmol, 8 pmol) or 2 pmol RPAP2-Pol II complex in a volume of 10 μl containing 30 mM HEPES pH 7.9, 100 mM KCl, 6 mM MgCl_2_, 2 mM DTT, 5% (v/v) glycerol for 30 min at 25°C. Reactions were initiated by the addition an equal volume of buffer containing 24 mM HEPES-KOH pH 8.0, 120 mM KCl, 10 mM MgCl_2_, 1.2 mM DTT, 24% (v/v) glycerol, 100 μg/ml BSA, 200 μM GTP, 200 μM CTP, 200 μM ATP, 200 μM UTP, and 99 nM [α-^32^P] UTP. The reactions were incubated at 25°C for 30 min and then were subjected to urea polyacrylamide gels and autoradiography.

### Generating dTAG endogenous knock-in and rescues cell lines

To generate RPAP2-dTAG cells by the endogenous knock-in, PITCh sgRNA/Cas9 and donor plasmids were mixed with 1×10^6^ DLD-1 cells followed by electroporation. After recovering for 2 days without antibiotic selection, cells were serially diluted and cultured with 1 µg/ml puromycin for 10-14 days. Single-clone colonies were picked, expanded, and genotyped by genomic DNA PCR targeting the integration site. For homogeneous knock-in clones, protein degradation efficiency was verified by DMSO and dTAG-13 treatment for 3 hours followed by western blotting.

To generate rescue cell lines, RPAP2-dTAG cells were initially infected with lentivirus expressing pLVX-Tet3G and cultured with neomycin for 2 weeks. Stable cells were transduced with lentivirus expressing wildtype RPAP2, RPAP2^Δ114-128^, or RPAP2^Y127A^ cloned into pLVX-Tet-On vector with Blasticidin resistance gene, and then selected with antibodies for 2 weeks.

### RNA interference

Lentivirus expressing short-hairpin RNAs was prepared by transfecting PLKO.1 shRNA plasmids and packaging plasmids containing psPAX2 and pMD2.G into 293T cells using PEI (Polysciences). Collected conditional media containing virus particles were used to transduce cells in the growth media supplemented with Polybrene for 24 hours. The infected cells were selected with 2 μg/ml puromycin for an extra 48 hours. The cells were then switched into growth media without antibiotics and grown for an additional 24 hours before being harvested for further analysis.

### ChIP-Rx and data analysis

ChIP-Rx was conducted following previously described (Orlando et al., 2014). Raw reads were trimmed by Trim Galore v0.6.6 (https://www.bioinformatics.babraham.ac.uk/projects/trim_galore/) to remove adaptors and low-quality sequences (-q 25) and then aligned to the human genome hg19 assembly using Bowtie2 v2.3.5.1 (Amemiya et al., 2019). PCR duplicates and low mapping quality reads (MAPQ < 30) were removed using Picard tools v2.23.3 (https://broadinstitute.github.io/picard/) (REMOVE_DUPLICATES = True) and SAMtools v1.9 (Li et al., 2009). Aligned read counts were then normalized to Reads Per Million mapped reads (RPM) using deeptools v3.5.0 (Ramirez et al., 2016),and blacklist regions for hg19 genome annotation from ENCODE project were removed (Amemiya et al., 2019). Reads aligned to human genome were normalized based on 1e6/ mm10_count calculated by SAMtools v1.9 (Li et al., 2009). Normalized bigwig files were generated by deeptools v3.5.0 (Ramirez et al., 2016).

### Identification of transcription start sites

To determine the genes expressed in DLD-1 cells, we used published PRO-cap bigwig file for DLD-1 cells to determine the transcription start site (TSS) (Aoi et al., 2020). RefSeq gene annotation was obtained from the UCSC Genome Browser and the transcription start sites were defined as the maximum PRO-cap signal site at the region between TSS –10 bp and TSS +300 bp. The transcript with maximum PRO-cap signal was selected as a representative gene for those protein coding genes with multiple isoforms.

**Figure S1.**
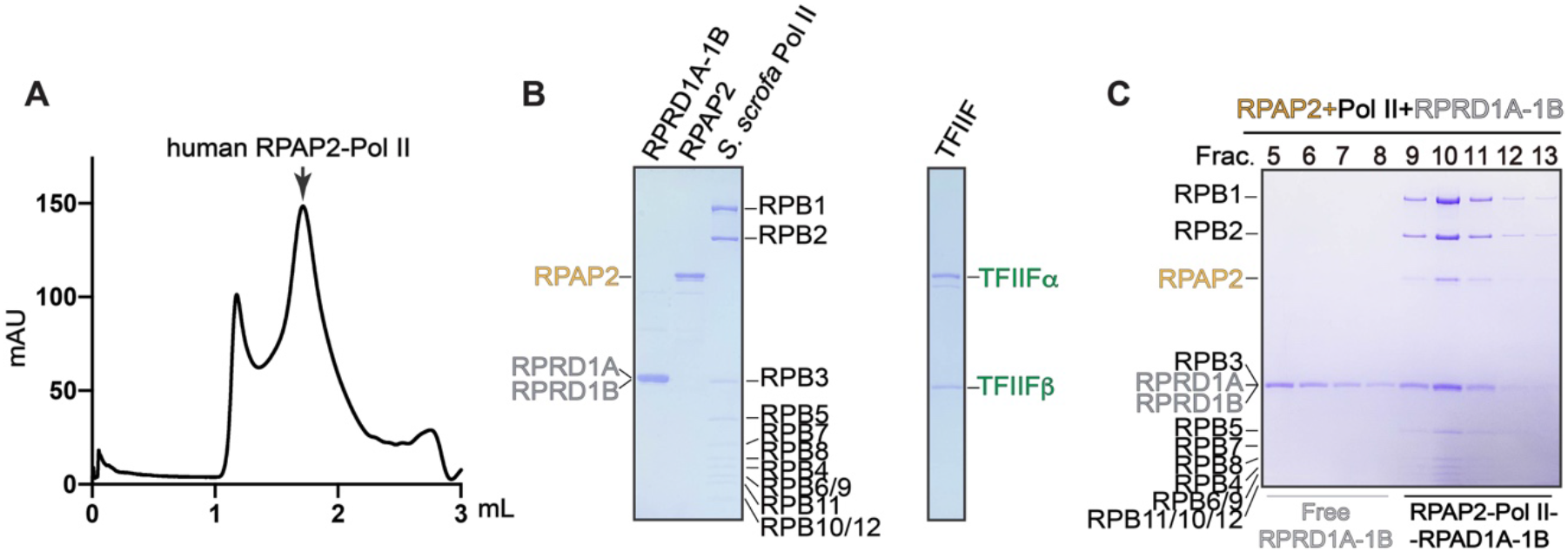
Preparation of RPAP2-Pol II complexes, Related to Figure 1 and 2. **(A)** Size exclusion chromatogram of the purified human RPAP2-Pol II complex. Pol II was overexpressed in Expi293F cells and endogenous RPAP2 was co-purified. **(B)** Purified human RPRD1A-RPAD1B, RPAP2, TFIIF and *S. scrofa* Pol II (4 residues substituted in H. sapiens Pol II) were subjected to SDS-PAGE and Coomassie blue staining. **(C)** Fractions of glycerol density gradient centrifugation of RPAP2-Pol II-RPRD1A-RPAD1B complex were subjected to SDS-PAGE followed by Coomassie blue staining.

**Figure S2.**
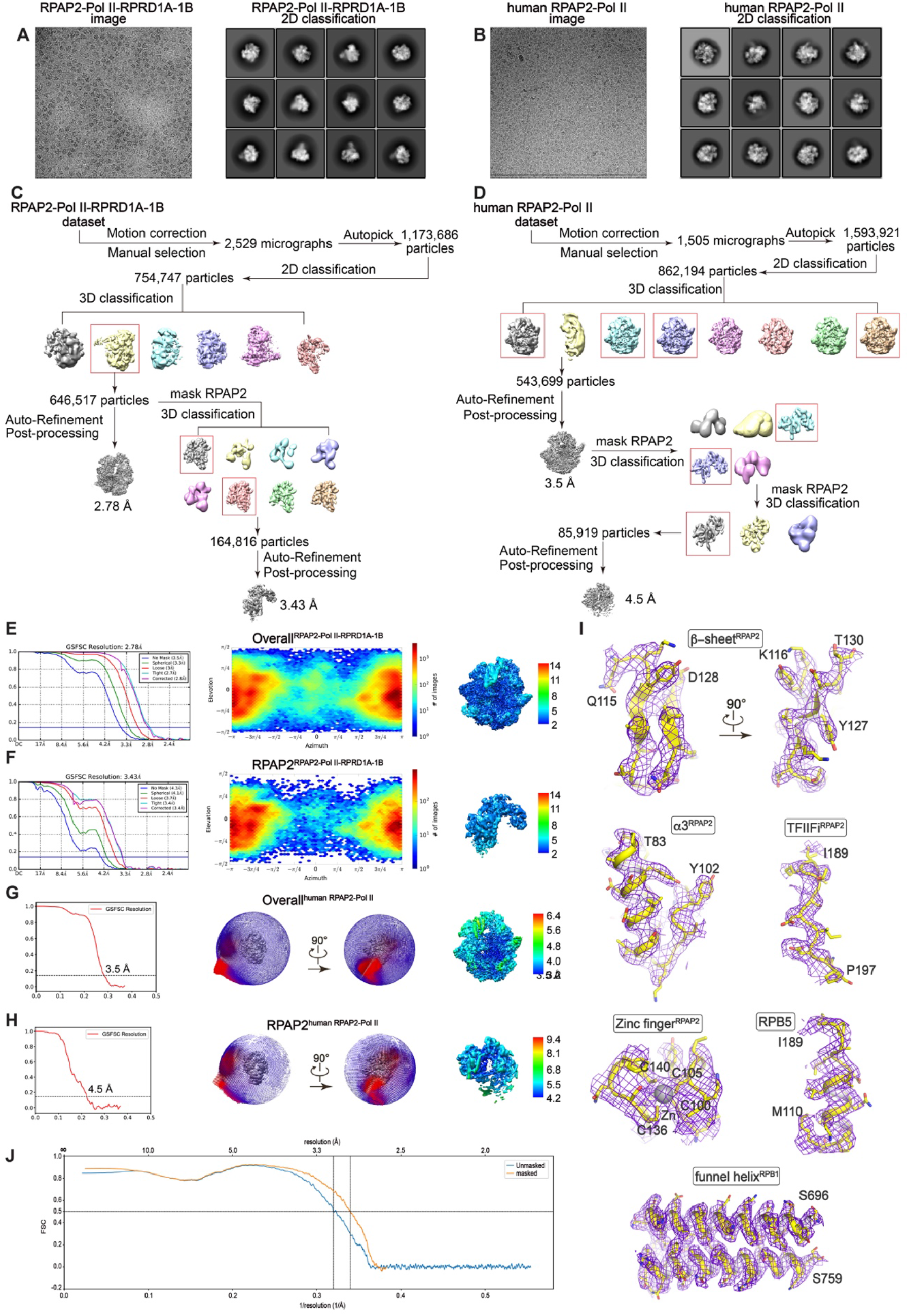
Structure determination and model building of RPAP2-Pol II complex, Related to Figure 2. (**A and B**) Representative cryo-EM raw micrographs (left panels) and 2D classification (right panels) of RPAP2-Pol II-RPRD1A-RPAD1B complex (A) and human RPAP2-Pol II complex (B). (**C and D**) Flow-charts of the cryo-EM image processing and 3D reconstruction for the RPAP2-Pol II-RPRD1A-RPAD1B complex (C) and human RPAP2-Pol II complex (D). (**E-H**) The GSFSC curves, angular distributions, and local resolution estimation of the cryo-EM reconstructions of RPAP2-Pol II-RPRD1A-RPAD1B complex (E and F) and human RPAP2-Pol II complex (G and H). **(I)** Representative structural models are shown with the corresponding cryo-EM maps shown in mesh. Residues are shown in sticks, indicating that the model was correctly built. **(J)** FSC curves between the model and cryo-EM map.

**Figure S3.**
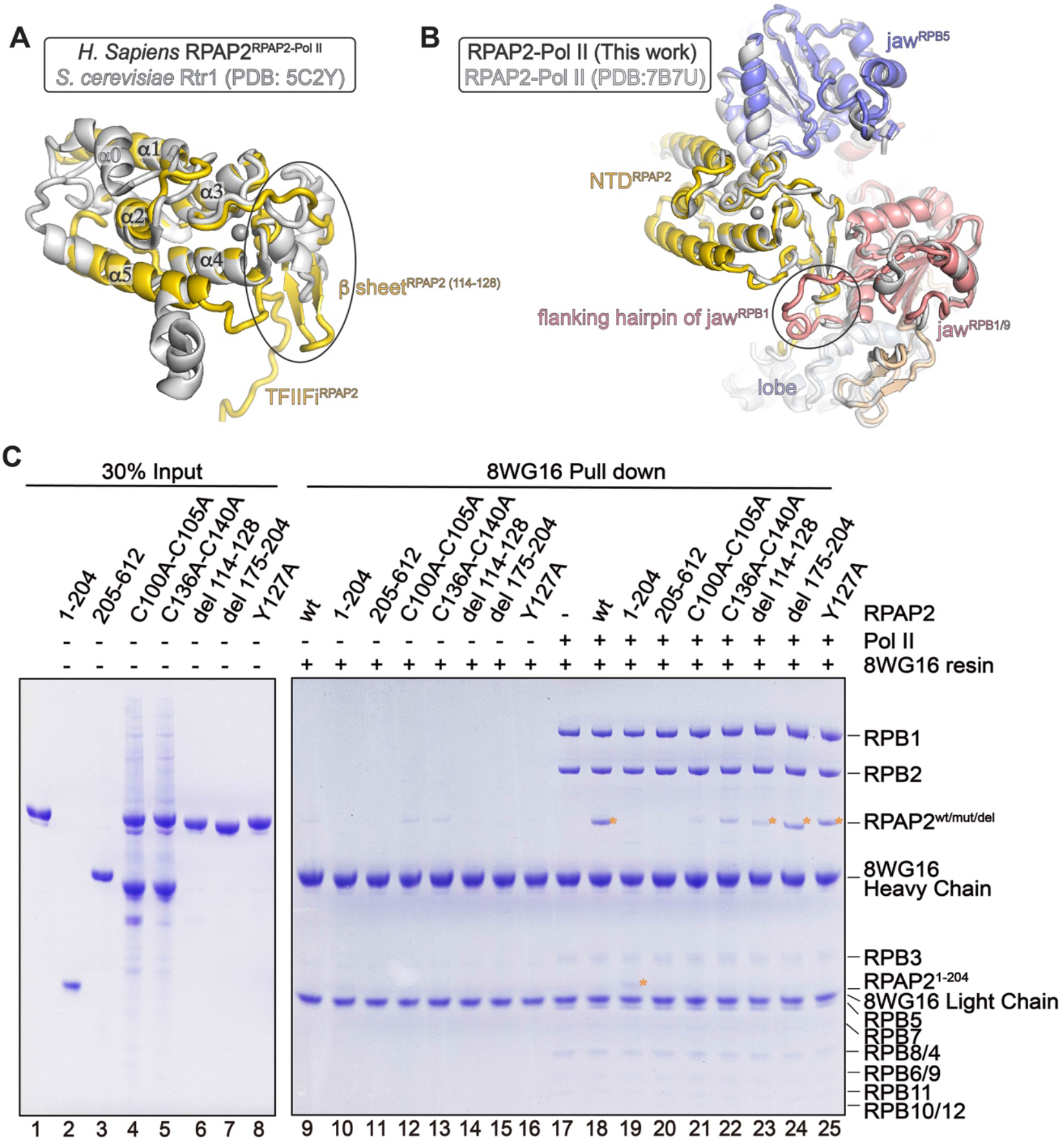
Interactions between RPAP2 and Pol II, Related to Figure 2. (**A and B**) Comparison of the structural models of RPAP2 (this work, colored) with isolated yeast Rtr1 (PDB: 5C2Y) (Irani et al., 2016) (B) and with human RPAP2-Pol II (gray, PDB: 7B7U) (Fianu et al., 2021). Structural differences are indicated with circles. (**C**) In vitro pull down using purified RPAP2 mutants/truncations and Pol II. The bound proteins were subjected to SDS-PAGE followed by Coomassie blue staining.

**Figure S4.**
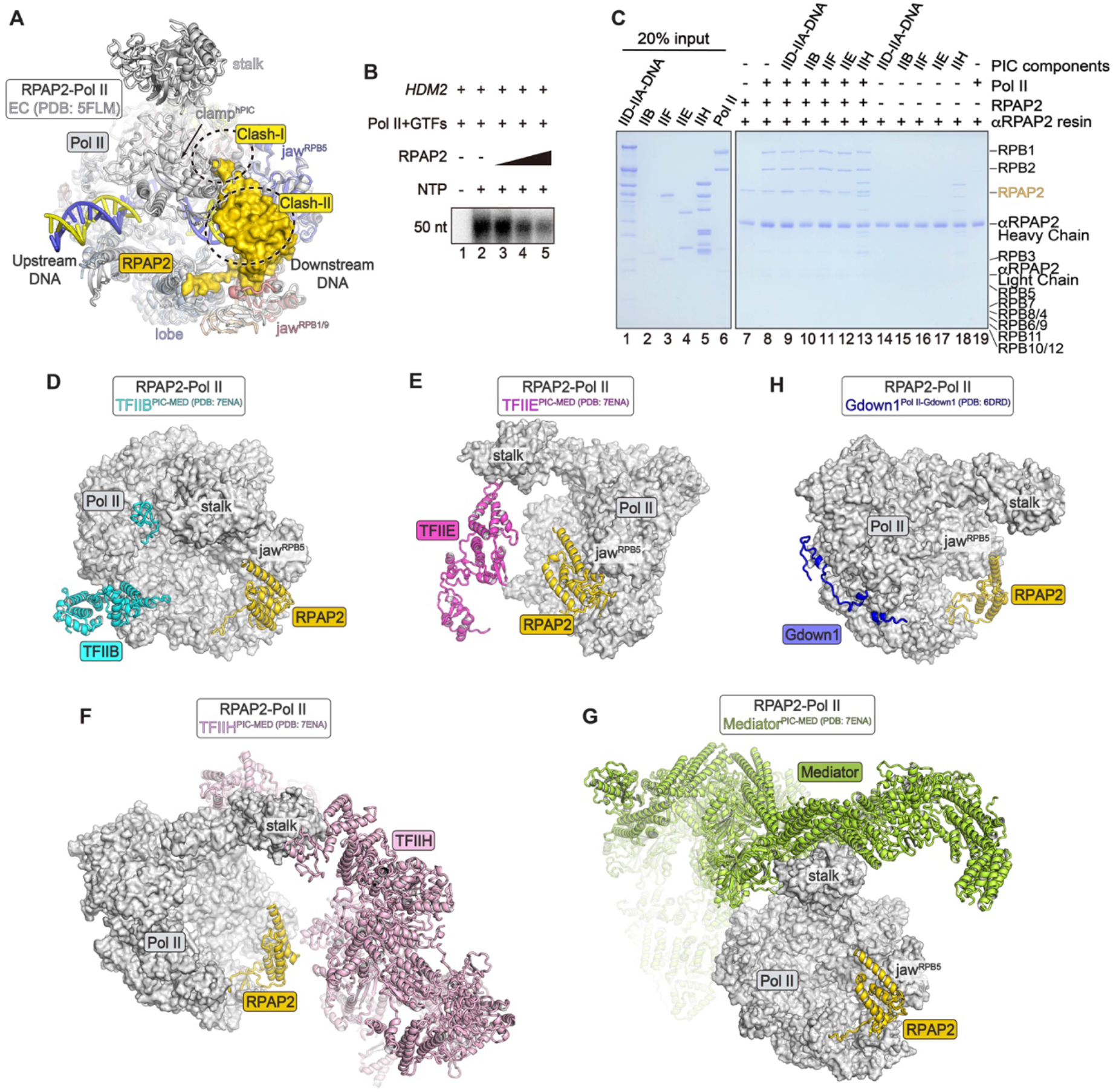
Steric clashes between RPAP2 and PIC/EC elements, Related to Figure 3. **(A)** Structures of RPAP2-Pol II and EC (PDB: 5FLM) (Bernecky et al., 2016) with Pol II superimposed. RPAP2-Pol II is colored as in Figure 2B and the EC is colored in gray, except that the DNA is colored in yellow and blue for clarity. **(B)** Autoradiogram of in vitro transcription initiation reactions. The complexes in the reactions included purified GTFs (TFIIA, TFIIB, TFIID, TFIIF, TFIIE and TFIIH), *HDM2* promoter, Pol II and an increasing amount of RPAP2. The reactions were performed essentially as Figure 3C except that RPAP2 was not pre-assembled with Pol II. **(C)** In vitro competitive binding assay using purified PIC components (TFIID-TFIIA-promoter, TFIIB, TFIIF, TFIIE and TFIIH), Pol II and RPAP2. RPAP2 and Pol II were pre-assembled and immobilized to the αRPAP2 resins followed by the addition of purified PIC components as indicated. The resins were extensively washed and subjected to SDS-PAGE and stained using Coomassie blue. (**D to H**) Position of RPAP2 on Pol II relative to other Pol II-binding proteins/complexes. Structure of RPAP2-Pol II was superimposed with structures of PIC-MED (PDB:7ENA) (Chen et al., 2021b) (C-F) and Pol II-Gdown1 (PDB: 6DRD) (Jishage et al., 2018) with Pol II shown in gray surface. TFIIB (D), TFIIE (E), TFIIH (F), Mediator (G), Gdown1 (H) and RPAP2 are shown as cartoon and the other GTFs were omitted for clarify. Structural comparison shows no steric clash between RPAP2 and these complexes.

**Figure S5.**
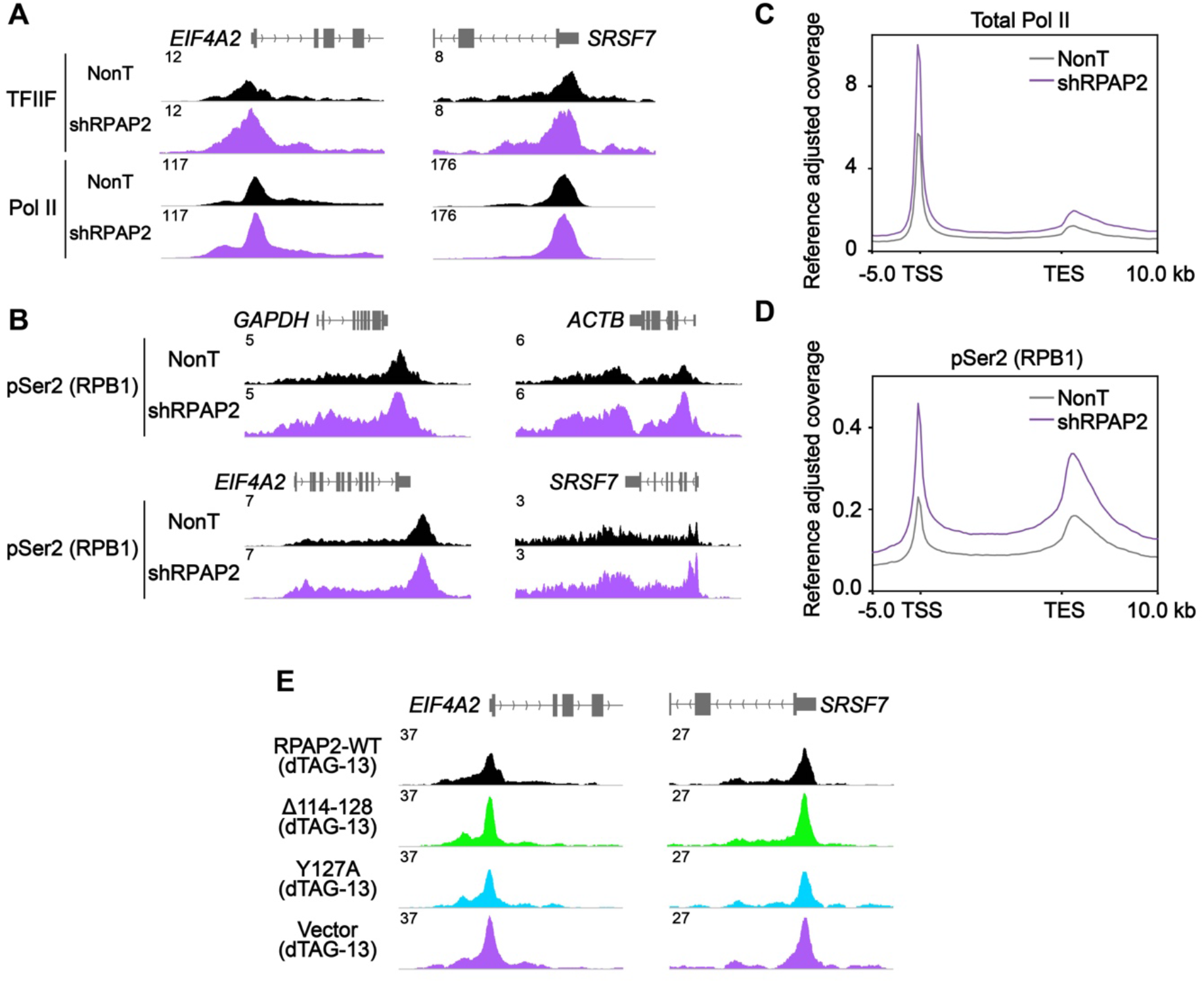
The impact of RPAP2 knockdown or degradation on PIC assembly at promoters, Related to Figure 4. **(A)** Representative track examples of genes *EIF4A2* and *SRSF7* showing the change of TFIIF and Pol II occupancy at promoters by RPAP2 knockdown. Pol II occupancy is represented by its largest subunit RPB1. **(B)** Representative track examples of genes *GAPDH, ACTB, EIF4A2* and *SRSF7* showing the change of Pol II pSer2 occupancy at gene bodies by RPAP2 knockdown. (**C and D**) Metaplot showing the occupancy of total Pol II (C) and pSer2 (D) at gene bodies measured by ChIP-Rx in NonT and RPAP2 knockdown cells. (**E**) Representative track examples of genes *EIF4A2* and *SRSF7* showing TFIIF occupancy in dTAG treated RPAP2-dTAG cells with induced expression of wildtype RPAP2, RPAP2^Δ114-128^, RPAP2^Y127A^, or vector.

**Table S1.**
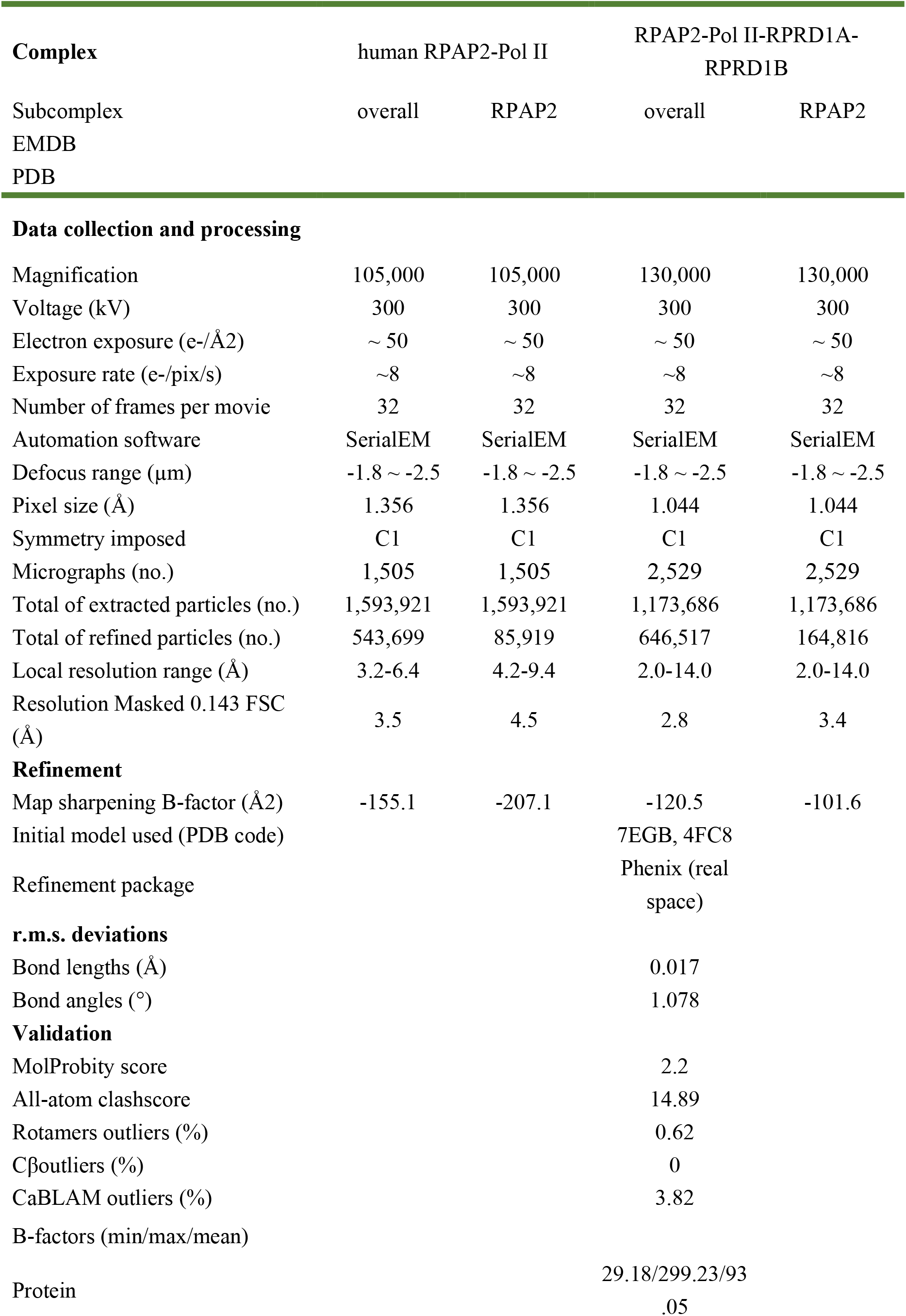
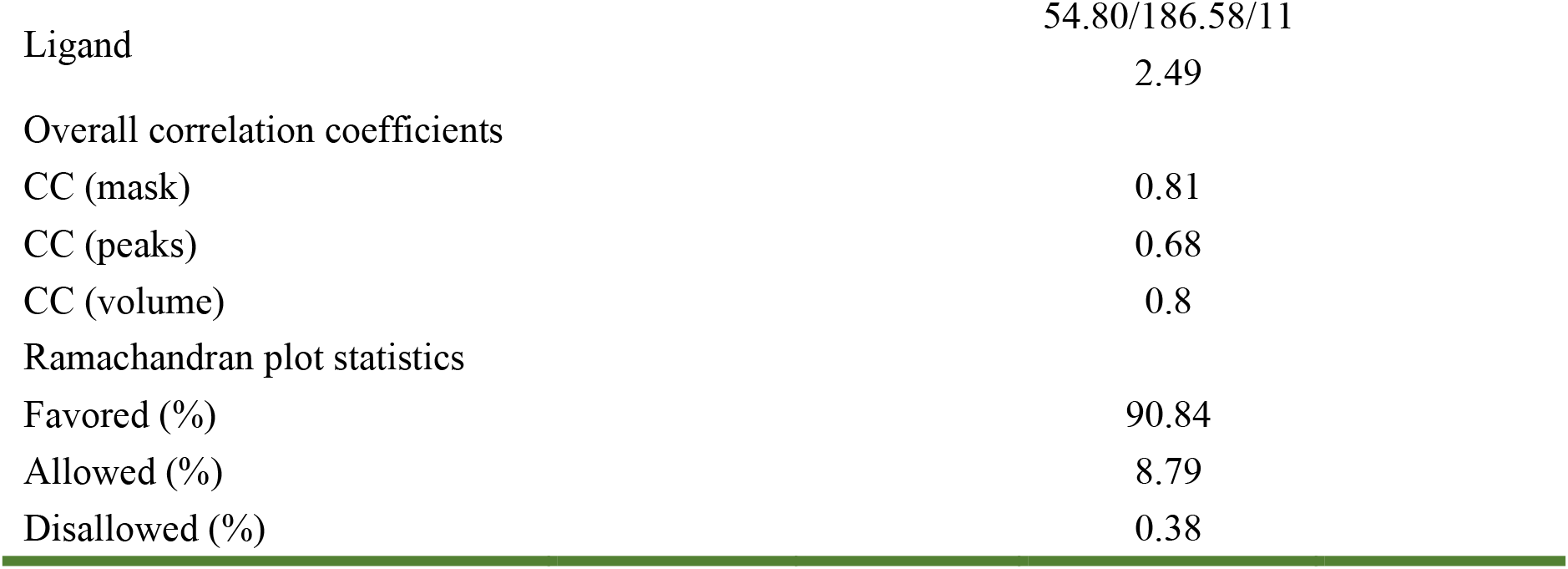
Cryo-EM data collection, refinement and validation statistics, Related to Figure2.

**Video S1. Overall structure of RPAP2-Pol II complex, Related to Figure2**.

The composite cryo-EM map and structural model of RPAP2-Pol II.

**Video S2. Steric clash between TFIIF and RPAP2, Related to Figure3**.

Structural comparison of RPAP2-Pol II with core PIC (cPIC) (PDB: 7EG7) (Chen et al., 2021a) with Pol II superimposed. Pol II is shown in surface and steric clash between TFIIF charge helix and TFIIFi^RPAP2^ is indicated with white circle. Unnecessary regions of cPIC were omitted for clarity.

